# Cortical tracking of a background speaker modulates the comprehension of a foreground speech signal

**DOI:** 10.1101/2021.04.12.439458

**Authors:** Mahmoud Keshavarzi, Enrico Varano, Tobias Reichenbach

## Abstract

Understanding speech in background noise is a difficult task. The tracking of speech rhythms such as the rate of syllables and words by cortical activity has emerged as a key neural mechanism for speech-in-noise comprehension. In particular, recent investigations have used transcranial alternating current stimulation (tACS) with the envelope of a speech signal to influence the cortical speech tracking, demonstrating that this type of stimulation modulates comprehension and therefore evidencing a functional role of the cortical tracking in speech processing. Cortical activity has been found to track the rhythms of a background speaker as well, but the functional significance of this neural response remains unclear. Here we employ a speech-comprehension task with a target speaker in the presence of a distractor voice to show that tACS with the speech envelope of the target voice as well as tACS with the envelope of the distractor speaker both modulate the comprehension of the target speech.

Because the envelope of the distractor speech does not carry information about the target speech stream, the modulation of speech comprehension through tACS with this envelope evidences that the cortical tracking of the background speaker affects the comprehension of the foreground speech signal. The phase dependency of the resulting modulation of speech comprehension is, however, opposite to that obtained from tACS with the envelope of the target speech signal. This suggests that the cortical tracking of the ignored speech stream and that of the attended speech stream may compete for neural resources.

**Significance Statement:** Loud environments such as busy pubs or restaurants can make conversation difficult. However, they also allow us to eavesdrop into other conversations that occur in the background. In particular, we often notice when somebody else mentions our name, even if we have not been listening to that person. However, the neural mechanisms by which background speech is processed remain poorly understood. Here we employ transcranial alternating current stimulation, a technique through which neural activity in the cerebral cortex can be influenced, to show that cortical responses to rhythms in the distractor speech modulate the comprehension of the target speaker. Our results evidence that the cortical tracking of background speech rhythms plays a functional role in speech processing.

## Introduction

Speech is a fascinatingly complex signal, the processing of which requires analysis of individual phonemes, syllables and words to extract meaning (Ingram, 2007; Poeppel *et al*., 2012; Hickok and Small, 2015). Moreover, when confronted with background noise such as other people talking, traffic or music, our brain must first segregate the target speech signal from the background acoustics before the processing of the target speech stream can begin (Bregman, 1994; Snyder and Alain, 2007).

An ignored speech signal can, however, still affect behaviour. Hearing one’s own name in the background can, for instance, shift our attention to the corresponding speaker (Moray, 1959; Wood and Cowan, 1995). Moreover, when attempting to listen to a target speaker in the presence of a distractor voice, a listener occasionally understands words from the distractor speaker (Brungart, 2001). A background speech signal in a listener’s native language is accordingly more distracting than speech in a foreign language or non-speech sounds such as music or traffic (Brungart, 2001; Cooke, Garcia Lecumberri and Barker, 2008; Cooke and Lu, 2010). However, the neural encoding of a background speech signal remains poorly understood.

An important aspect of speech processing in a computer is the parsing of the speech stream into functional constituents such as syllables and words. The brain presumably employs a similar parsing. Cortical activity, in particular in the delta (1 - 4 Hz) and theta (4 - 8 Hz) frequency ranges, has indeed been found to track speech rhythms, such as those set by the rate of syllables and words (Giraud and Poeppel, 2012; Ding and Simon, 2014; Di Liberto, O’Sullivan and Lalor, 2015). A computational model shows that such tracking can yield effective online syllable parsing (Hyafil *et al*., 2015).

Cortical tracking of speech rhythms may contribute to the processing of both an attended and an ignored speech signal. When selectively attending to one of two competing speakers, cortical activity tracks the rhythms of the attended speech as well as those of the ignored voice, although the cortical tracking of the attended speaker is stronger than that of the unattended speaker (Ding and Simon, 2012; Mesgarani and Chang, 2012; Horton, D’Zmura and Srinivasan, 2013). Moreover, the cortical tracking of an attended talker is modulated by speech comprehension, such as when speech is noise-vocoded, presented in different levels of background noise or compared between a native and foreign language (Ding, Chatterjee and Simon, 2014; Brodbeck, Hong and Simon, 2018; Broderick *et al*., 2018; Vanthornhout *et al*., 2018; Etard and Reichenbach, 2019; Weissbart, Kandylaki and Reichenbach, 2019).

The modulation of the cortical speech tracking by cognitive factors such as attention and comprehension are not merely side effects of other neural processes. Instead, a functional role of the cortical tracking of an attended speaker in speech processing has been shown through transcranial alternating current stimulation (tACS), a non-invasive technique to influence cortical activity in the frequency range of the stimulation (Antal and Paulus, 2013; Helfrich *et al*., 2014). Applying tACS with waveforms that were derived from the speech envelope, reflecting the speech rhythms, has demonstrated an effect on the cortical speech tracking (Zoefel, Archer-Boyd and Davis, 2018). Moreover, this type of tACS has been found to modulate the comprehension of speech in noise, as well as the comprehension of noise-vocoded speech (Riecke *et al*., 2018; Wilsch *et al*., 2018; Kadir *et al*., 2020; Keshavarzi *et al*., 2020a).

However, it remains unclear what functional role the cortical tracking of an ignored speech signal has, and whether it can influence behaviour. Here we therefore set out to investigate the impact of tACS with the speech envelope of a distractor voice on speech comprehension.

## Materials and Methods

### Experimental design and statistical analysis

To investigate the functional role of the cortical tracking of a background speech signal, we employed a simplified version of a cocktail party with two voices, a male and a female one (Figure 1A,B). Subjects were instructed to listen to a particular target speaker and to ignore the other distractor voice. Their auditory cortices were stimulated simultaneously and noninvasively with alternating current (Figure 1A). The current waveforms were derived from the speech envelope of either the target or the distractor speech signal (Figure 1B).

**Figure 1:**
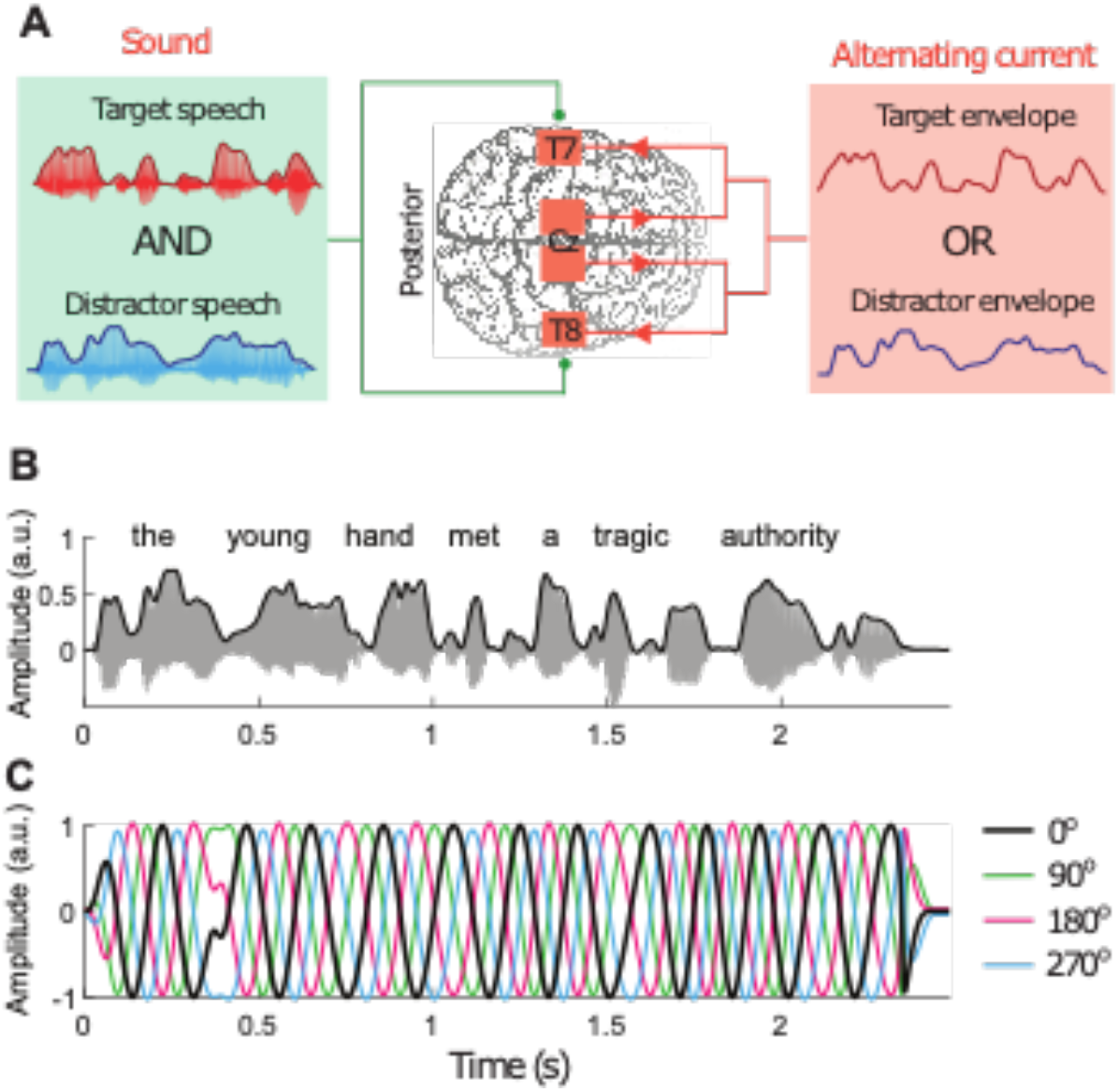
Experimental setup. (**A**), Subjects listened to two competing voices. They were instructed to attend to one of them that served as the target while the other represented the distractor. Subjects simultaneously received tACS derived from the envelope of either the target or of the distractor speech stream. (**B**), Both target and distractor speech were single, semantically unpredictable sentences (acoustic waveform, grey; envelope, black). (**C**), The current waveforms were obtained by band-pass filtering the speech envelope in the theta frequency range and by processing the signal so that all maxima and minima occurred at equal values. The obtained signal was then shifted by four different phases (0°, black; 90°, green; 180°, red; 270°, blue). Note that the signal shifted by 180° (red) is indeed the inverse of the signal without phase shift (black).

Most of the power spectral density of the speech envelope is contained in two frequency bands, the delta band (1 - 4 Hz) and the theta band (4 - 8 Hz). Previously we showed that the modulation of speech comprehension results from stimulation with the theta frequency portion but not from the delta frequency portion of the speech envelope (Keshavarzi *et al*., 2020a), and that speech comprehension can be enhanced when the current has no temporal delay with respect to the speech stream (Keshavarzi and Reichenbach, 2020). In this study we accordingly employed current waveforms that represented the theta-band portion of the speech envelope and that were temporally aligned to the acoustic signal (Figure 1C).

We determined the modulation of speech comprehension through tACS by shifting the current waveforms by four different phases: 0°, 90°, 180°, and 270°. We previously showed that tACS with the theta-band portion of the envelope of a target speech then leads to a modulation of speech comprehension that depends on the phase shifts in a cyclical manner (Keshavarzi *et al*., 2020a).

To assess the potential dependence of the speech comprehension scores *CS*(*ϕ*_*n*_) on the phase shifts *ϕ*_*n*_ = *n·* 90° with *n* = 0,1,2,3, we first determined, for each subject, the mean comprehension score 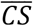 across the different phase shifts,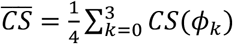. We then focussed on the differences of the speech comprehension scores Δ*CS*(*ϕ*_*k*_) of each subject from the mean value, 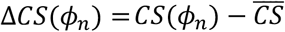. Due to the cyclical nature of the phase shifts, the dependence of this change in the speech comprehension score on the phase shifts *ϕ*_*n*_ could be expressed through the Discrete Fourier Transform as

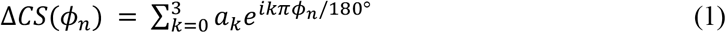

in which the complex coefficients *a*_*n*_ were calculated as 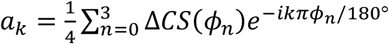. Because the speech comprehension scores were real, it followed that the coefficients *a*_0_ and *a*_2_ were real as well and that 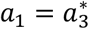. Equation (1) for the dependence of Δ*CS*(*ϕ*_1_) on the phases *ϕ*_*n*_ could thus be rewritten as

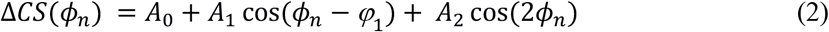

in which *A*_0_*= a*_0_ was a constant offset, *A*_1_ = 2 |*a*_1_| and *A*_2_*= a*_2_ respectively denoted the amplitude of the variation at the periods of 360° and 180°, and *φ*_1_ = *arg(a*_1_) was the phase offset at the first period.

To fit Equation (2) to the data, we first rewrote the middle term on the right-hand side, *A*_1_ cos(*ϕ*_*n*_ − *φ*_1_), as a sum of a sine and a cosine function of *ϕ*_*n*_ to obtain

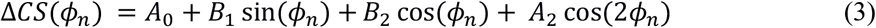

with the coefficients *B*_1_*= A*_1_ cos(*φ*_1_) and *B*_1_*= A*_1_ *sin*(*φ*_1_). We then determined the parameters *A*_0_, *B*_1_, *B*_2_, and *A*_2_ through multiple linear regression. The corresponding *p-*values were corrected for multiple comparisons using the FDR correction. The amplitude *A*_1_ of the variation at the period of 360° followed as 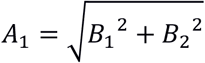 and the corresponding phase offset as *φ*_1_ = arctan(*B*_2_*/B*_1_).

Because the data may contain outliers, we also employed robust regression to determine the parameters *A*_0_, *B*_1_, *B*_2_, and *A*_2_ in Equation (3) (Welsch, 1977). Robust regression employs an iteratively reweighted least-squares algorithm that assigns weights to each data point. Outlier data points thereby obtain a lower weight, such that they contribute less to the parameter estimation.

To determine whether the robust regression yielded a better fit than the standard regression, we employed bootstrapping with 10,000 samples. For each sample, we performed both the standard regression as well as the robust regression, and computed the resulting *r*^2^ value for each method. We then compared the two distributions of *r*^2^ values through a paired one-tailed Student’s t test.

To investigate the subject-to-subject variability of speech comprehension for tACS with both target and distractor envelopes, we determined the phases that, for each type of stimulation and for each individual subject, yielded the highest comprehension score. For every subject we then aligned the phase with respect to this best phase. This left us with three remaining phase shifts that were measured relative to the best phase. The dependence of the changes in the speech comprehension scores, Δ*CS*, on these three relative phase shifts *ϕ*_*n*_*=n·* 90° with *n=* 1,2,3 could be described by

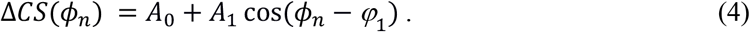

This equation could be recast as

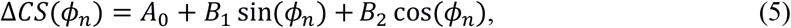

in which the coefficients *B*_1_ and *B*_2_ followed as *B*_1_*= A*_1_ cos(*φ*_1_) and *B*_2_*= A*_2_ cos(*φ*_1_). We determined the parameters *A*_0_, *B*_1_ and *B*. through multiple linear regression, and corrected the resulting *p*-values for multiple comparisons through the FDR correction. As for the case when the phase was not aligned to the best phase, the amplitude *A*_1_ of the variation at the period of 360° in Equation (4) could be computed as 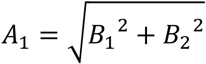, and the phase offset as *φ*_1_ = arctan(*B*_2_*/B*_1_).

To account for potential outliers, we also determined the parameters *A*_0_, *B*_1_ and *B*_2_ in Equation (4) through robust regression, that is, using the iteratively reweighted least-squares algorithm. We then assessed whether the robust regression yielded a better fit than the standard regression by bootstrapping as described above.

### Participants

Eighteen native English speakers, eight of them female, with self-reported normal hearing participated in the experiment. The number of participants was chosen based on previous experiments on tACS with the speech envelope, including our own, which revealed that such a number of participants is sufficient to yield effects on speech comprehension on the population level (Riecke *et al*., 2018; Wilsch *et al*., 2018; Kadir *et al*., 2020; Keshavarzi *et al*., 2020; Keshavarzi and Reichenbach, 2020).

The participants were between 18 and 29 years, with a mean age of 23.8 years. All participants were right-handed and had no history of mental health problems or neurological disorders. Before starting the experiment, participants gave informed consent. The experimental protocol was approved by the Imperial College Research Ethics Committee.

### Data and code availability

Data and code are available upon request.

### Hardware set-up

The acoustic stimuli as well as the current signals were presented using a PC with a Windows 7 operating system. Both types of signals were synchronized on the PC, and then converted to analogue signals using a USB-6212 BNC device (National Instruments, U.S.A.). The current signals were fed into a splitter connected to two neurostimulation devices (DC-Stimulator Plus, NeuroConn, Germany). The acoustic stimuli were passed through a soundcard (Fireface 802, RME, Germany) connected to earphones (ER-2, Etymotic Research, U.S.A.).

### Acoustic stimuli

We employed semantically unpredictable sentences, e.g. “The charitable stresses think the unusual investigation.”. These were created in text form using the Python package Natural Language Toolkit (Bird, Klein and Loper, 2009; Beysolow II, 2018). The texts were then converted to sound through the software TextAloud, using either a female or a male voice, with a sampling rate of 44.1 kHz. Each sentence consisted of seven words that included five key words; these were used to determine the subject’s comprehension score. The target sentences were presented at a SPL level of 60 dB SPL.

### tACS waveforms

We employed nine different types of tACS waveforms. One waveform was a sham stimulus that consisted of a short current that started at the beginning of the target speech and lasted 500 ms. Smooth onsets and offsets were implemented through ramps with a duration of 100 ms. Four further waveforms were computed from the envelope of the target speech by band-pass filtering this signal in the theta frequency band (zero phase IIR filter, low cutoff of 4 Hz, high cutoff of 8 Hz, order 6). The band-pass filtered envelopes were further processed so that all maxima and minima occurred at equal values, and were then shifted by four different phases (0°, 90°, 180°, 270°). The four remaining types of tACS waveforms were computed from the envelope of the distractor speech; these computations were analogous to the computation of the waveforms from the envelope of the target speech.

### Experimental procedure

Participants were seated in a soundproof room. Two rubber electrodes (35 cm^2^) covered by sponge pads and wetted by a 0.9% saline solution (5 ml per electrode) were then placed on the T7 and T8 locations and two further ones to the left and right of the Cz location of the participant’s head (International 10-10 system). The electrode placed on T7 and the one to the left of Cz were connected to one neurostimulation device and the two remaining ones to another device. The electrodes at the T7 and T8 locations functioned as the anodes and the ones at Cz as the cathodes. The resistance between electrodes of each device was set to be almost equal and below 10 kΩ.

To determine the maximum magnitude of current stimulation for each subject, a five-second sinusoidal signal with a frequency of 3 Hz was presented to her or him. The signal amplitude was initially 0.1 mA and was increased in steps of 0.1 mA up to a maximum of 1.5 mA. The increase in amplitude was stopped once the subject reported a skin sensation, and the amplitude used in the prior step was employed as the maximum magnitude of current stimulation during the next parts of the study. Across the participants we thereby obtained a mean amplitude of 0.67 mA with a standard deviation of 0.24 mA.

The participants listened to diotically-presented acoustic stimuli through earphones. For each participant, we determined the signal-to-noise ratio (SNR) corresponding to the comprehension level of 50% using an adaptive procedure (Kollmeier, Gilkey and Sieben, 1988; Kaernbach, 2001) while the sham stimulation was applied. To this end, the subject was asked to listen to the male voice and ignore a competing female voice, and to then repeat what they heard. The procedure was started with a randomly selected SNR between –3 dB and 0 dB. If the participant understood no more than two keywords correctly, the SNR was then increased by 1 dB for the next trial; otherwise, it was decreased by 1 dB. The procedure was stopped after seven reversals in the SNR or after running 17 trials. This was done four times for each participant. The average of the last three SNR values during the last three repetitions was considered as the 50% sentence reception threshold (SRT). Across subjects we thereby obtained a mean SRT of –6.9 dB and a standard deviation of 1.6 dB. To save measurement time, the SRT was only determined for the male voice as the target speaker, and not also for the female voice as the target of attention. Although the SRT for the female voice might differ slightly from the SRT for the male speaker, such differences could not bias the subsequent results which were based on the comprehension of both the male and the female voice taken together.

For each subject we then employed the subject-specific SRT to measure their speech comprehension under tACS with the speech envelope. For each type of tACS waveform, we presented subjects with 26 sentences in noise. 13 of these 26 sentences had the female voice as the target and a male voice as distracting talker, and the other 13 sentences *vice versa*. The relative contribution of the male and of the female voice in the acoustic mixture therefore depended on whether the male or the female voice was the target of attention. In particular, the SRT was chosen per subject as determined from the adaptive procedure. Because the SRTs were negative for all subjects, this meant that, when attention was directed towards the male voice, the male voice was fainter than the female voice. On the other hand, when the subject was instructed to attend the female voice, the female voice had an amplitude below that of the male voice.

Both target and distracting sentences were randomly selected in each trial and were unknown to both the experimenter and the subject. The order of attention was randomly determined, and subjects were informed over a display before each sentence whether they should attend to the male or the female speaker. After listening to a sentence, the participant repeated what they understood. The response was recorded by a microphone. The recorded responses were manually graded by the experimenter to determine the percentage of correctly understood words. The participant took a brief rest for three minutes after every 60 trials.

## Results

We first verified that the envelopes of the target and of the distractor sentences were indeed unrelated. We found that Pearson’s correlation coefficient between the two envelopes was –0.003 ± 0.002 (mean and standard error of the mean, average over 100 pairs of sentences). The correlation coefficient was therefore small and not significantly different from zero (*p* = 0.98, two-tailed Student’s t-test).

We then proceeded to analyze the effect of tASC on speech comprehension. We found that tACS with the target envelope resulted indeed in a significant modulation (Figure 2A, Table 1). Both standard multiple linear regression and robust multiple linear regression to fit Equation (3) to the data showed a dependence of the speech comprehension score on the phase shift of the stimulation. In particular, the parameter *B*_2_ was significantly different from 0, although the parameters *B*_1_ and *A*_2_ Were not. The dependence of the speech comprehension scores on the phase shift of the stimulation waveform emerged therefore at the longest possible period of 360°, but not at the shorter period of 180°.

**Table 1:**
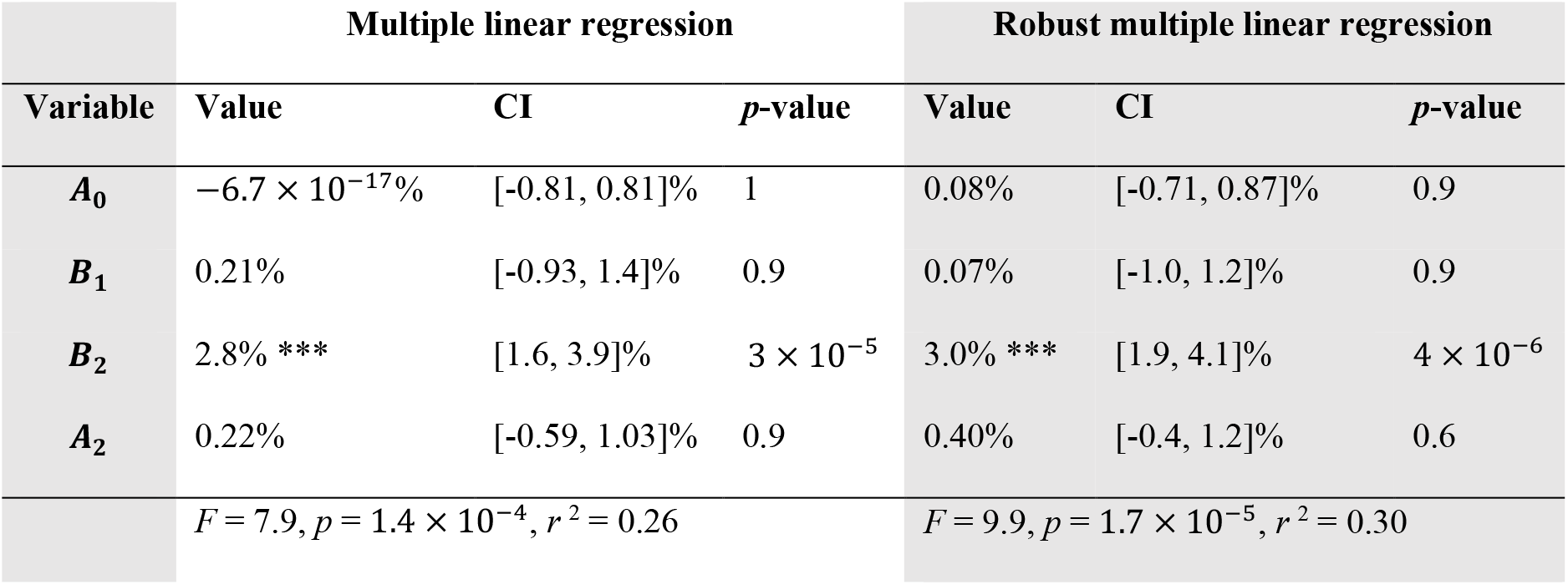
Multiple linear regression for the dependence of the speech comprehension scores on the stimulation phase, for stimulation with the target envelope. Left: results from multiple linear regression using the least square estimate. Right: results from robust multiple linear regression, using an iteratively reweighted least squares algorithm. The values for the parameters as well as for the confidence intervals (CI) are given in %. The *p*-values have been corrected for multiple comparisons through the FDR correction. Statistical significance is denoted by asterisks (*, 0.01 < *p* ≤ 0.05; **, 0.001 < *p* ≤ 0.01; ***, *p* ≤ 0.001).

**Figure 2:**
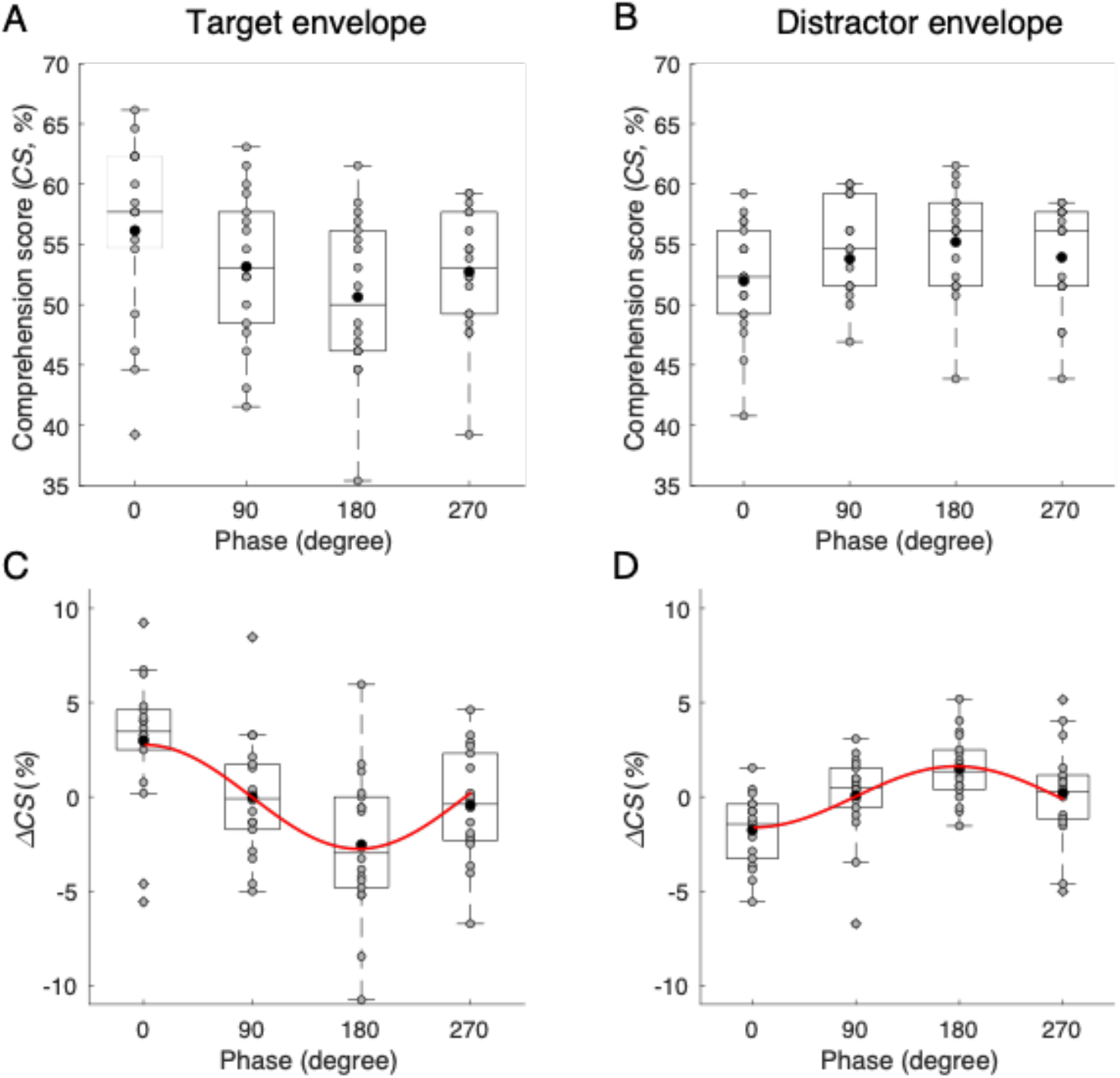
Modulation of speech comprehension through tACS. (**A, B**), Speech comprehension scores (CS) at different phase shifts of the current waveform are shown as box plots. In addition, the scores from each subject are shown as grey disks; black disks denote the mean values. (**C, D**), The difference Δ*CS* of the comprehension score to the mean score per subject across the four phases. The red curves show the fit obtained from equation (3) through robust regression, including only the terms that are statistically significant. (**C**), tACS with the target envelope yielded the highest speech comprehension at a phase shift of 0°, and the lowest at the phase shift of 180°. (**D**), tACS with the distractor envelope led to a significant modulation of speech comprehension as well, although with a different phase dependence: the highest score occurred at the phase shift of 180° and the lowest at the phase shift of 0°.

Because of the insignificance of the parameter *B*_1_, the amplitude *A*_1_ of the variation at the period of 360° followed simply as the parameter *B*_2_: *A*_1_ = *B*_2_. The corresponding phase shift was *φ*_1_ = 0°.

Robust regression yielded a significantly higher *r* ^2^ value than the standard regression (*p* < 2× 10^−10^, bootstrap with 10,000 samples and paired one-tailed Student’s t test). Stimulation with tACS with the target envelope thus modulated speech comprehension by about 3.0%, the value of *A*_1_ = *B*_2_ obtained from the robust regression.

Our results regarding the dependence of speech comprehension on the phase of tACS with the target envelope largely replicated our earlier findings (Keshavarzi *et al*., 2020a). Indeed, we previously showed that this type of tACS yields a modulation of speech comprehension such that the highest comprehension score is observed at 0° phase shift, and the lowest at a phase shift of about 240°. Here we observed similarly that no phase shift yielded the maximal speech comprehension, while we obtained the lowest comprehension scores at a phase shift of 180°. The moderate deviation in the phase shift of the lowest comprehension score might be explained by the different number of phase shifts that we employed in the two studies, as well as by experimental uncertainties.

Armed with the ability to assess the influence of tACS on speech comprehension, we then analyzed the effect of tACS with the distractor envelope (Figure 2D, Table 2). As before, we employed multiple linear regression with FDR correction for multiple comparisons to fit Equation (3) to the data and to analyze the statistical significance of the different terms (Figure 2, Table 2). As for tACS with the target envelope, we found that the parameter *B*_2_ was significant, but not the parameters *B*_1_ and *A*_2_. Importantly, tACS with the distractor envelope did therefore lead to a significant modulation of speech comprehension.

**Table 2:** Multiple linear regression for the dependence of the speech comprehension scores on the stimulation phase, for stimulation with the distractor envelope. The presentation of the results is as in Table 1.

| Variable | Multiple linear regression |  |  | Robust multiple linear regression |  |  |
| --- | --- | --- | --- | --- | --- | --- |
| | Value | CI | $p$-value | Value | CI | $p$-value |
| $A_0$ | $3.5 \times 10^{-16}\%$ | $[-0.5, 0.5]\%$ | 1 | 0.13% | $[-0.34, 0.61]$ | 0.8 |
| $B_1$ | -0.06% | $[-0.77, 0.65]\%$ | 1 | 0.10% | $[-0.57, 0.77]\%$ | 0.8 |
| $B_2$ | -1.6% *** | $[-2.3, -0.9]\%$ | $9 \times 10^{-5}$ | -1.6% *** | $[-2.2, -0.9]\%$ | $6 \times 10^{-5}$ |
| $A_2$ | -0.14% | $[-0.64, 0.36]\%$ | 1 | -0.26% | $[-0.74, 0.21]\%$ | 0.5 |
| | $F = 7.0, p = 3.4 \times 10^{-4}, r^2 = 0.20$ | | | $F = 7.8, p = 1.5 \times 10^{-4}, r^2 = 0.26$ | | |

The modulation emerged at the period of 360°. Because of the insignificance of the parameter *B*_1_, and the negativity of the parameter *B*_2_, the amplitude *A*_1_ followed as *A*_1_ = −*B*_2_, and the corresponding phase shift as *φ*_1_ = 180°. The amplitude of the modulation of speech comprehension through stimulation with tACS with the distractor envelope was therefore about 1.6%. As for stimulation with the target envelope, we found that the robust regression gave a significantly better fit to the data than the standard regression (*p* < 2× 10^−10^, bootstrap with 10,000 samples and paired one-tailed Student’s t test).

Although tACS with the target envelope and tACS with the distractor envelope both modulated speech comprehension at the longest-possible period of 360°, there were two important differences between the two types of stimulation. First, the phase shifts that led to the best and worst speech comprehension scores were the opposite in the two cases. While tACS with the speech envelope at a phase shift of 0° led to the best speech comprehension when based on the envelope of the target speech, this phase shift produced the worst speech comprehension when derived from the envelope of the distractor speech. Inversely, a phase shift of 180° gave the lowest comprehension scores for tACS with the target envelope, but the highest comprehension score for tACS with the distractor envelope.

Second, the strength of the modulation of speech comprehension differed between tACS with the target envelope and tACS with the distractor envelope. Specifically, considering the results from the robust regression, tACS with the target envelope yielded a modulation amplitude of *A*_1_ = 3% that was higher than the modulation amplitude of *A*_1_ = 1.6% that we obtained from tACS with the distractor envelope (*p* = 0.037, two-tailed Student’s t-test). We note, however, that the *p*-value was close to 0.05, and that the difference between the amplitudes *A*_1_ became insignificant when considering the results from the standard regression (*p* = 0.1, two-tailed Student’s t-test).

Previous research investigated phase shifts between tACS with the speech envelope and the acoustic waveform in the case of rhythmic speech, where syllables occurred at a specific and fixed frequency. These studies found that the tACS did modulate speech comprehension, but that the best and worst phase shifts were highly variable between subjects (Riecke *et al*., 2018; Zoefel, Archer-Boyd and Davis, 2018). Similarly, related work on temporal delays between the sound and the neurostimulation waveform found large variability between subjects, and in one case no significant effect on the population level (Riecke *et al*., 2018; Wilsch *et al*., 2018; Erkens *et al*., 2020; Wang *et al*., 2020). In our previous work on the role of phase shifts in tACS with the speech envelope we employed, as in this study, natural speech in which syllables did not occur at a single fixed frequency and found a phase dependency of the resulting speech comprehension that was largely consistent across the different subjects.

We therefore sought to investigate the subject-to-subject variability of speech comprehension for tACS with both target and distractor envelopes. To this end, we first determined for every individual subject and for each of the two stimulation types the phase that yielded the highest comprehension score. We then measured the phase relative to this best phase for that particular subject (Figure 3A,B).

**Figure 3:**
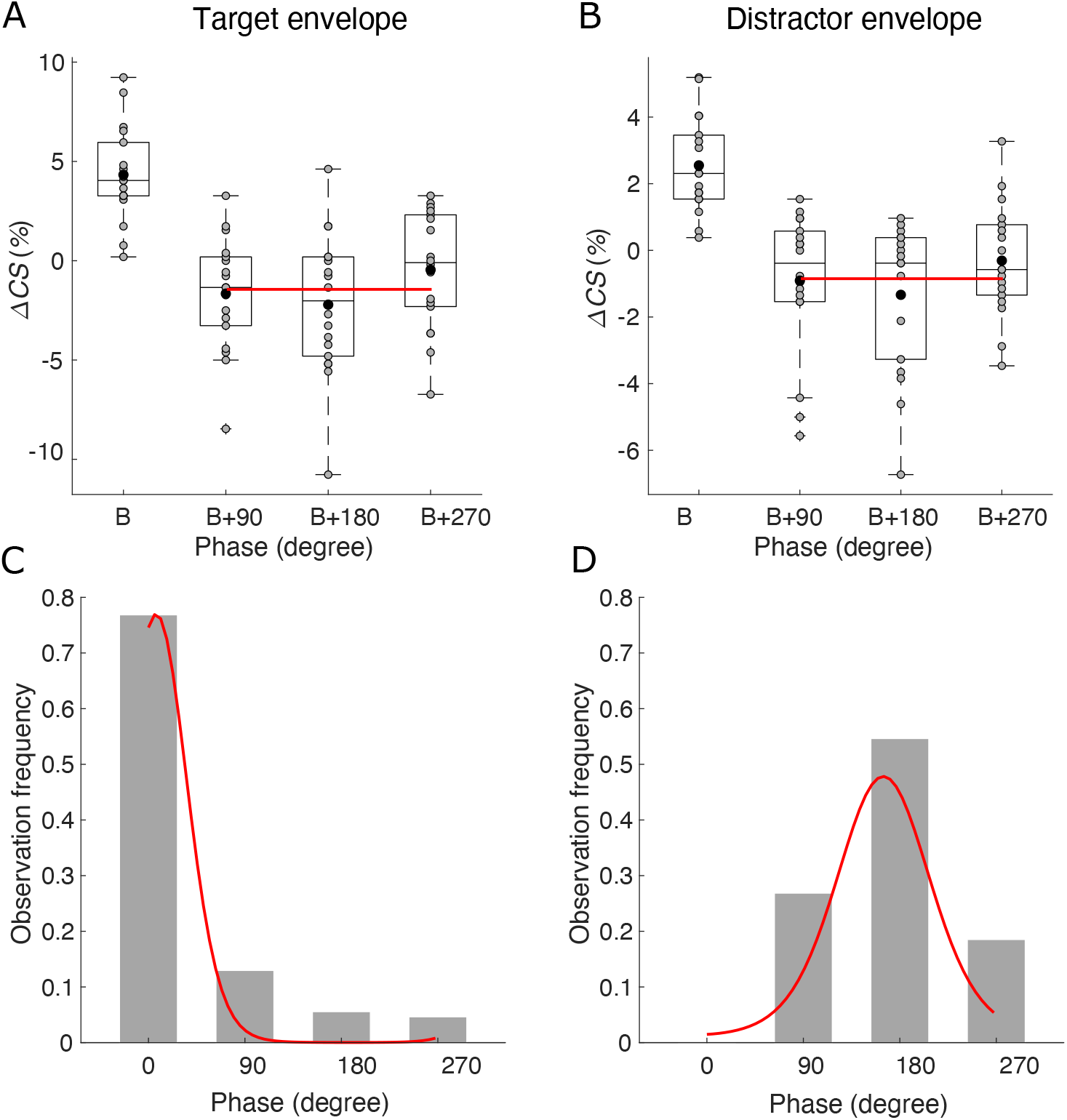
Consistent phase dependency across subjects. **(A, B)**, After aligning the phase relative to the best phase (‘B’) per subject, neither tACS with the target envelope (**A**) nor tACS with the distractor envelope (**B**) lead to a significant effect of the relative phase on speech comprehension. Grey disks show the scores from individual subjects, and the black disks denote the population mean. The red line denotes the fit obtained from the model described by equation (5) with only the significant terms included. **(C, D)**, The distributions of the best phases of the different subjects were significantly different from a uniform distribution, and could be fitted well by von Mises distributions (red lines). (**C**), For tACS with the target envelope, the best phases occurred mostly around 0°. (**D**), In contrast, tACS with the distractor envelope yielded best phases that clustered around 180°.

We employed multiple linear regression as well as robust regression to fit Equation (5) to the data and to determine the statistical significance of the parameters. We found, however, no significant modulation of speech comprehension by the relative phases, neither for tACS with the target envelope (Table 3) nor for tACS with the distractor envelope (Table 4). These results showed that the alignment to the best phase increased rather than decreased the noise in the data, indicating that the best phase did not vary substantially between participants.

**Table 3:** Multiple linear regression for the dependence of the speech comprehension scores on the stimulation phase, aligned for the best phase and for stimulation with the target envelope. The presentation of the results is as in Table 1.

| Variable | Multiple linear regression |  |  | Robust multiple linear regression |  |  |
| --- | --- | --- | --- | --- | --- | --- |
|  | Value | CI | *p*-value | Value | CI | *p*-value |
| $A_0$ | -1.06% | [-2.11, -0.01]% | 0.14 | -0.96% | [-2.04, 0.11]% | 0.23 |
| $B_1$ | 0.59% | [-1.65, 0.45]% | 0.26 | -0.60% | [-1.68, 0.47]% | 0.26 |
| $B_2$ | 1.15% | [-0.66, 2.97]% | 0.26 | 1.13% | [-0.73, 2.99]% | 0.26 |
| | $F = 1.47, p = 0.24, r^2 = 0.0544$ | | | $F = 1.41, p = 0.25, r^2 = 0.0523$ | | |

**Table 4:** Multiple linear regression for the dependence of the speech comprehension scores on the stimulation phase, aligned for the best phase and for stimulation with the distractor envelope. The presentation of the results is as in Table 1.

| Variable | Multiple linear regression |  |  | Robust multiple linear regression |  |  |
| --- | --- | --- | --- | --- | --- | --- |
|  | Value | CI | *p*-value | Value | CI | *p*-value |
| $A_0$ | -0.60% | [-1.28, 0.07]% | 0.23 | -0.44% | [-1.11, 0.22]% | 0.56 |
| $B_1$ | -0.30% | [-0.97, 0.37]% | 0.37 | -0.11% | [-0.78, 0.56]% | 0.74 |
| $B_2$ | 0.72% | [-0.44, 1.89]% | 0.33 | 0.52% | [-0.64, 1.67]% | 0.56 |
| | $F = 1.18, p = 0.32, r^2 = 0.0442$ | | | $F = 0.841, p = 0.44, r^2 = 0.0319$ | | |

To further investigate the inter-subject variability of the modulation of speech comprehension through tACS, we computed histograms of the best phases of each subject (Figure 3C,D). For tACS with the target envelope, we found that the distribution of the best phases was significantly different from a uniform distribution (*p* = 2 × 10^−1^, Rayleigh test). The distribution had a maximum at a phase shift of 0° and a mean phase shift of 4.4°, with an angular deviation of 0.74 and a resultant vector length of 0.72. The distribution could be fitted well with the unimodal von Mises distribution (*p* = 0.005, Watson’s test). This was in line with our finding above that the phase shift of 0° led to the highest speech comprehension scores for this type of tACS (Figure 2A).

The distribution of the best phases for tACS with the distractor envelope was significantly non-uniform as well (*p* = 0.002, Rayleigh test). However, the best phases occurred around phase shifts of 180°: the maximum of the distribution was at 180° and its mean was 169°, with an angular deviation of 0.93 and a resultant vector length of 0.57. The unimodal nature of the distribution was confirmed by fitting a von Mises distribution (*p* = 0.005, Watson’s test). The peak in the distribution coincided with the maximum of the speech comprehension scores at this phase shift (Figure 2B).

Although the angular deviation of the distribution for the best phases obtained for tACS with the distractor envelope was larger than the one for tACS with the target envelope, the difference was not significant (*p* = 0.32, concentration homogeneity test).

We also investigated whether tACS with the target or distractor envelope could improve speech comprehension beyond sham stimulation. We therefore compared the speech comprehension scores at the best phase shift of 0° for tACS with the target envelope, as well as at the best phase shift of 180° for tACS with the distractor envelope, to the scores under sham stimulation (Figure 4). We found that there was no statistically significant difference between the different comprehension scores (one-way ANOVA, *df* = 2, *F* = 1.72, *p* = 0.2).

**Figure 4:**
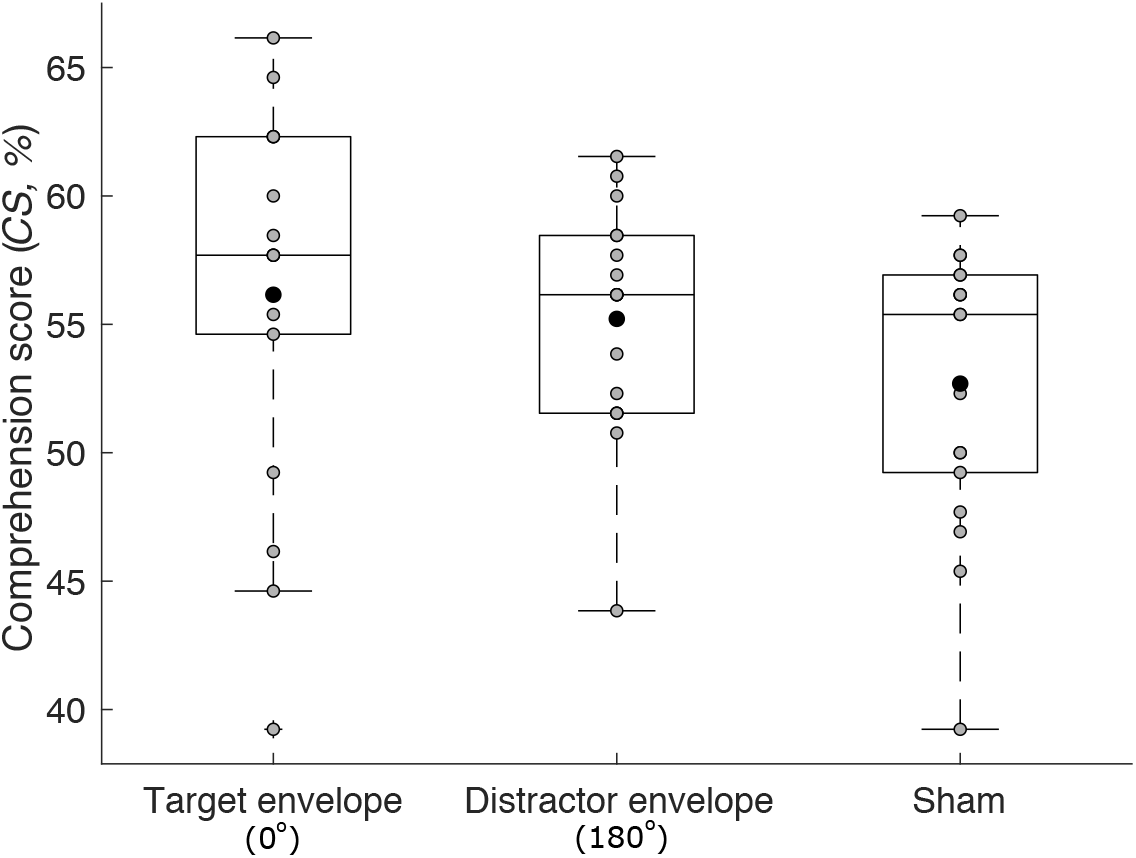
Comparison to sham stimulation. Because we obtained the highest speech comprehension for tACS with the target envelope at a phase shift of 0°, and for tACS with the distractor envelope at a phase shift of 180°, we compared the speech comprehension scores for these two stimulation types to sham stimulation (box plots; grey disks, individual subjects; black disks, mean scores). The speech comprehension scores showed no significant difference between the three conditions.

## Discussion

Taken together, our results showed that tACS with the distractor envelope influenced speech comprehension, evidencing that the cortical tracking of the ignored speaker plays a functional role in speech processing. Indeed, because the target and the distractor speech signals that we employed were unrelated, the envelope of the distractor speech carried no information about the target signal. tACS with the distractor envelope could not therefore influence the cortical tracking of the target speech, but only that of the distractor speech.

tACS with the target envelope influenced speech comprehension in a very similar manner to tACS with the distractor envelope. In particular, both types of stimulation led to a modulation of speech comprehension that varied sinusoidally, at the longest possible period, with the applied phase shifts. Moreover, both types of tACS led to a modulation of speech comprehension that had a largely consistent phase dependency across the different subjects. In particular, while earlier studies showed that stimulation parameters such as delay or phase shift could influence speech comprehension in a manner that was highly variable from subject to subject (Riecke *et al*., 2018; Wilsch *et al*., 2018; Zoefel, Archer-Boyd and Davis, 2018), our results did not show such variability. Indeed, when adjusting the phase shift to the ‘best’ value per subject, we no longer obtained a significant modulation of speech comprehension, indicating that the adjustment increased rather than decreased the noise in the data. This result was in line with our previous studies on the effects of tACS with the speech envelope on speech comprehension (Kadir *et al*., 2020; Keshavarzi and Reichenbach, 2020; Keshavarzi *et al*., 2020b).

Despite these similarities between tACS with the target and with the distractor envelope, there were two major differences. First, the phase shift that led to the highest speech comprehension score was 0° for tACS with the target envelope and 180° for tACS with the distractor envelope. This difference suggests that tACS with the envelope of a speech signal may act on the early perceptual separability of that signal from an acoustic background. At a phase shift of 0°, tACS with a speech envelope appeared to enhance the neural representation of that speech stream. tACS with a speech envelope shifted by 180°, in contrast, presumably led to a suppression of the neural encoding of the corresponding speech stream.

This interpretation suggests that the cortical tracking of a speech signal plays, at least partly, a role in auditory stream formation, both for an attended and for an ignored acoustic source (Petersen *et al*., 2017; Hausfeld *et al*., 2018; Fiedler *et al*., 2019). Our finding that tACS with the distractor envelope modulates the comprehension of the target speech then implies that the formation of the two auditory streams, of the target and the distractor speech, are not independent of each other. Instead, an enhanced representation of one acoustic stream appears to cause a reduced representation of the other stream. Further investigation of such a competition between the cortical tracking of the foreground and of the background acoustic signals may employ neuroimaging to quantify the neural responses, for instance paired to tACS and behavioural assessments as explored here.

As the second major difference between tACS with the target and with the distractor envelope, the former type of stimulation led to a stronger modulation of speech comprehension than the latter type. The difference in the modulation amplitudes emerged when analyzing the data with robust regression, accounting for outliers in the data, but not when employing standard regression. We found indeed that robust regression gave significantly better fits to the data than standard regression. The resulting higher certainty in the fit parameters obtained from robust regression as compared to standard regression presumably allowed to obtain a statistically-significant difference in the modulation of speech comprehension between stimulation with the target and with the distractor envelope.

This stronger modulation may point to an effect of tACS with the target envelope that goes beyond auditory stream formation, such as by aiding the parsing of the attended speech stream into syllables and words. Previous research has indeed reported that the comprehension of noise-vocoded speech, in the absence of background noise, can be modulated by tACS with the speech envelope, evidencing that this type of tACS can influence speech processing beyond acoustic stream formation (Riecke *et al*., 2018). Because we found that the modulation of speech comprehension through tACS with the target envelope was about 3.0%, almost twice the corresponding value of 1.6% found for tACS with the distractor envelope, our results suggest that tACS with the envelope of a speech signal influenced predominantly auditory stream formation, but also had a sizable impact on the further neural processing of that speech stream.

Our results therefore suggest that cortical tracking of an ignored speaker plays a functional role in auditory stream formation, while the cortical tracking of an attended speaker likely goes beyond that to encompass further semantic and syntactic processing. These findings are in line with recent results on the encoding of multiple talkers in the auditory cortex. Cortical responses in Heschel’s gyrus have indeed been found to be able to encode an ignored speaker, whereas cortical activity in the superior temporal gyrus responds more to an attended talker (O’Sullivan *et al*., 2019). In addition, early cortical tracking of acoustic onsets in mixtures of speech signals can occur even for onsets of an ignored speaker that are masked by acoustic activity of a target speaker, suggesting that the activity of the auditory cortex can recover such masked ignored signals, while later activity is restricted to features of an attended speaker (Brodbeck *et al*., 2020). Furthermore, cortical tracking of speech rhythms has been shown to encode information on the lexical as well as on the semantic level for an attended speech stream, but not for an ignored one (Brodbeck, Hong and Simon, 2018; Broderick *et al*., 2018). In addition, behavioural studies have found accordingly that listeners can notice simple acoustic changes in an ignored voice but fail to understand it (Cherry, 1953; Deutsch and Deutsch, 1963; Treisman, 1964; Kidd *et al*., 2005). However, we also note that speech can be more distracting when intelligible, suggesting that some linguistic information is processed even in ignored speech (Dai *et al*., 2017).

We found previously that tACS with the target envelope could improve the comprehension of speech in speech-shaped noise beyond sham stimulation (Keshavarzi and Reichenbach, 2020; Keshavarzi *et al*., 2020a). In our present study, however, the speech comprehension scores obtained during tACS with the ‘best’ phase shifts were not significantly higher than those obtained under sham stimulation. This might be due to the different background noise that we used here, namely a competing talker rather than the previously-employed speech-shaped noise. The competing talker provided informational masking, while speech-shaped noise contributed only energetic masking (Pollack, 1975; Kidd *et al*., 2008). Informational and energetic masking affect speech comprehension differently (Brungart, 2001; Freyman, Balakrishnan and Helfer, 2004; Kidd *et al*., 2016). Moreover, most of the recent studies on tACS with the speech envelope found a detrimental rather than beneficial impact of the stimulation on speech comprehension (Riecke *et al*., 2018; Wilsch *et al*., 2018; Kadir *et al*., 2020). In addition, as opposed to speech-shaped noise, the amplitude of the competing talker fluctuated in time, allowing for dip listening (Miller and Licklider, 1950; Bacon and Wesley Grantham, 1989; Gustafsson and Arlinger, 1994; Rosen *et al*., 2013). Both factors may influence how tACS with the speech envelope affects speech comprehension. Disentangling these aspects will further clarify how cortical tracking of speech rhythms can impact speech processing, and potentially lead to the development of optimized tACS waveforms for speech-in-noise enhancement.

## Acknowledgement

This research was supported by the Royal British Legion Centre for Blast Injury Studies, by EPSRC grants EP/M026728/1 and EP/R032602/1, as well as by the U.S. Army through project 71931-LS-INT.

